# Vision-Language Models for Image-Based Dietary Assessment: A Benchmark of Accuracy, Cost, and Prompt Strategies Across Ten Models

**DOI:** 10.64898/2026.07.26.740845

**Authors:** Sam Sterling, Lauren T. Berube, Andrea J. Glenn, Aasma Shaukat, Souptik Barua, Morgan E. Grams, Aristotelis Tsirigos

## Abstract

**Background:** Dietary assessment is the cornerstone of clinical management and research studies evaluating diet and health. Traditional methods such as food diaries and 24-hour recalls can be burdensome, prone to recall bias, and difficult to adhere to. Image-based dietary assessment using vision-language models (VLMs) offers a potential solution.

**Objective:** Our goal was to benchmark state-of-the-art VLMs for automated food recognition, weight estimation, and calorie estimation using Google’s Nutrition5k dataset.

**Methods:** We evaluated 3,229 food images using ten approaches: proprietary VLMs (Gemini 2.0 Flash, 2.5 Flash, 3.0 Flash, and 3.1 Flash Lite; GPT- 4o, GPT-4o-mini, and GPT-5 Mini; and Claude Haiku 4.5), an open-source VLM (Qwen2-VL-7B), and a commercial food recognition API (FatSecret). We assessed calorie and weight estimation using Lin’s Concordance Correlation Coefficient (CCC) and component detection using Jaccard similarity.

**Results:** Gemini 3.0 Flash achieved the best calorie estimation (CCC 0.767, MAE 80.7 kcal), while Gemini 3.1 Flash Lite offered very comparable accuracy (CCC 0.754) with the highest ingredient recognition (Jaccard 0.655) at the lowest cost among top-performing models ($0.59/1K images). Among earlier-generation models, Gemini 2.0 Flash remained competitive (CCC 0.742, Jaccard 0.621) at a fraction of the cost ($0.10/1K images). A human validation study in which four annotators reviewed 440 images revealed systematic omissions in the original Nutrition5k labels. After correction, the extrapolated ingredient-overlap score for Gemini 2.0 Flash increased from 0.62 to an estimated 0.82, suggesting that raw Jaccard scores substantially underestimate true model performance.

**Conclusions:** Current VLMs can perform automated dietary assessment with reasonable accuracy from single overhead photographs. Our results inform model selection for dietary assessment applications and highlight remaining challenges in calorie estimation and component detection for complex, multi-item meals.

## Introduction

Dietary intake assessment remains a challenge in clinical settings. Traditional methods, including 24-hour dietary recalls, food frequency questionnaires, and written food diaries, can be prone to biases including recall bias, portion size misestimation, and ingredient omission. These tools can also be tedious and labor intensive to complete, and require a certain level of literacy. These limitations have motivated research into automated image-based dietary assessment. [1–3] At the same time, the consumer nutrition tracking market has experienced rapid growth over the last five years and is projected to reach USD 4.56 billion by 2030. [4] Partially as a result of this growth, the past decade has seen substantial investment in the use of machine learning for food image analysis. While early approaches employed convolutional neural networks for meal classification tasks, [5–6] recent and more sophisticated systems combine food recognition with portion estimation, but require specialized depth sensors, reference objects, or multi-view capture to estimate food volumes. [7–11] However, this is not how food is typically logged for consumer health applications; meals are usually recorded as an overhead smartphone photo taken under variable conditions, making accurate weight and calorie estimation challenging. As a result, most commercially available nutrition apps identify foods in an image, estimate a plausible portion size or weight, and then match those items to entries in a nutrition database to return calories and nutrients. [6] Several AI-powered nutrition apps, such as Keenoa [12] and Rx Food, [13] have undergone validation studies and can return macronutrient and micronutrient information by linking recognized foods to curated nutrient databases. However, these commercial solutions use proprietary recognition pipelines that cannot be independently benchmarked or customized by researchers. In contrast, general-purpose vision-language models (VLMs) are accessible via pay-per-use APIs at transparent and often lower cost [14], require no specialized infrastructure beyond an API key, and can be adapted to specific research needs through prompt engineering, which we show can meaningfully improve accuracy.

Recent studies have explored the use of publicly available VLMs for nutrition estimation with promising initial results. [15–16] By pre-training on internet-scale image-text pairs, these models are able to acquire visual understanding capabilities that transfer to specialized domains without task-specific fine-tuning. [17–23] For dietary assessment, VLMs offer potential advantages: they can recognize diverse food items through general object understanding, leverage nutritional knowledge embedded in their text training, and provide structured outputs for downstream processing.

Our goal was to benchmark state-of-the-art VLMs for automated food recognition, as well as weight and calorie estimation, using Google’s Nutrition5k dataset. By conducting a systematic comparison of proprietary VLMs, open-source architectures, and specialized commercial APIs, this study establishes a performance benchmark to guide model selection and deployment in real-world dietary assessment applications.

## Methods

In this work, we used Google’s Nutrition5k dataset, which consists of 5,000 human-annotated overhead photographs of real meals served at Google’s cafeteria intended to capture the natural variability of real-world meals. We evaluated ten approaches: proprietary VLMs (four Gemini models, three GPT models, and Claude Haiku 4.5), an open-source VLM (Qwen2-VL-7B), and a commercial food recognition API (FatSecret). Using Gemini 2.0 Flash, we also tested five prompt variants.

### Dataset and Preprocessing

Google’s Nutrition5k dataset [24] consists of 5,000 unique cafeteria meals collected with a custom scanning rig at Google cafeterias. Each meal represents a unique tray and may contain a single food item or multiple components; the dataset does not include beverages. For each meal, Nutrition5k provides a single top-down (overhead) red-green-blue (RGB) image; many meals also include a registered overhead depth map and four rotating side-view videos. Specific collection dates are not reported in the Nutrition5k documentation. Each meal includes dish-level totals for mass, calories, and macronutrients derived from the dataset’s measurement pipeline, as well as a component list that we use only for ingredient-overlap evaluation (this list can be coarser than what is visually identifiable in the image). For our test, we used only the overhead RGB photograph available in the realsense_overhead directory, and we applied the following inclusion criteria: (1) not marked as deprecated in metadata, (2) valid calorie and weight values (non-zero, non-missing), and (3) at least one labeled component. Of the 5,000 meals in Nutrition5k, 3,490 had overhead RGB images available. We excluded 31 meals containing deprecated ingredients, 2 plate-only entries, and 228 with zero or missing weight values, yielding a final sample of 3,229 images.

### Models Evaluated

We evaluated ten models spanning proprietary VLMs, an open-source VLM, and a commercial food recognition API. From Google, we tested four Gemini models: 2.0 Flash, 2.5 Flash, 3.0 Flash, and 3.1 Flash Lite. From OpenAI, we tested GPT-4o, GPT-4o-mini, and GPT-5 Mini. We also evaluated Anthropic’s Claude Haiku 4.5. As an open-source representative, we included Alibaba’s Qwen2-VL-7B-Instruct [25] (hereafter Qwen2-VL-7B), which we ran locally on our high-performance computing (HPC) cluster using NVIDIA A100 GPUs. Lastly, we included the leading commercial food database and nutrition recognition API, FatSecret. [26] Our prompt engineering study was conducted using Gemini 2.0 Flash only (the most cost-effective of the commercial VLMs at the time of study); the best-performing prompt (P3) was then applied to all models for the full benchmark. FatSecret does not accept custom prompts, so it was evaluated using its default pipeline. Each model’s response was parsed as structured JSON; responses that could not be parsed into the expected schema were excluded from analysis without retries. All performance metrics were computed over successfully parsed responses only.

### Prompt Engineering Study

All VLMs except FatSecret received a standardized prompt requesting structured JavaScript Object Notation (JSON) output with the following fields:

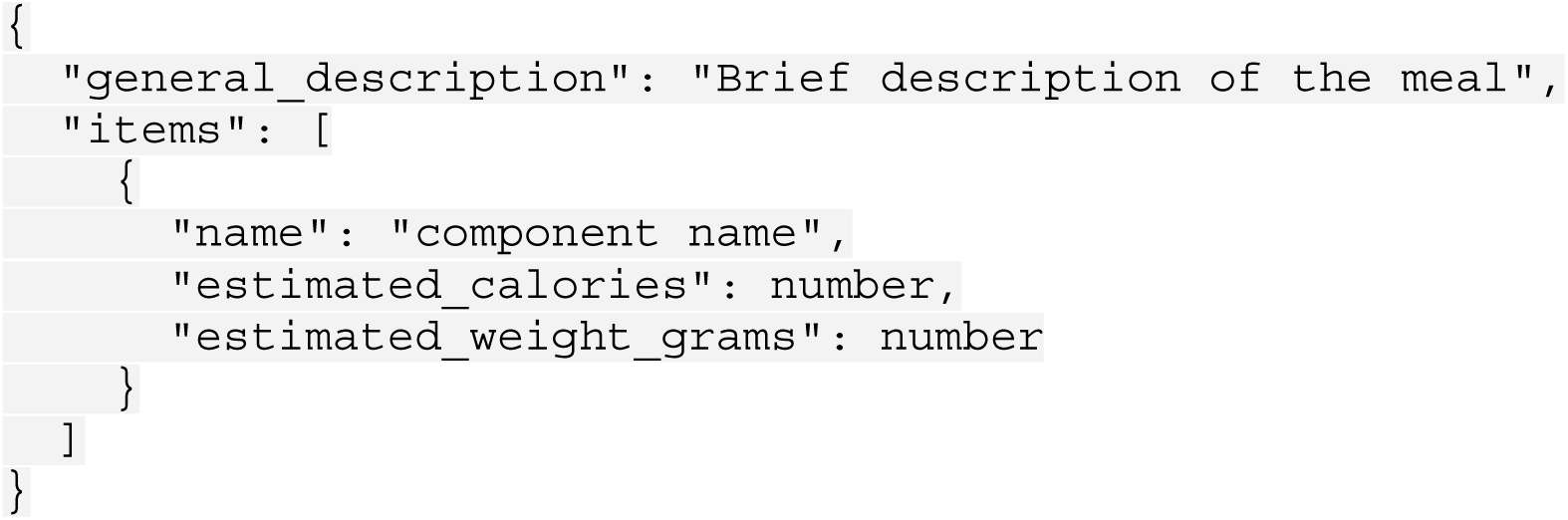

We developed and evaluated five prompt variants (P1 through P5), as prompting has been shown to materially affect reasoning in large language models. [27–29] Our prompt engineering study was conducted using Gemini 2.0 Flash, the cheapest commercial VLM in our study. Temperature was set to 0.1 for all models across both the prompt ablation and full benchmark to encourage deterministic outputs; other generation parameters (top_p, max_tokens) were left at provider defaults.

We first evaluated four single-pass prompt variants (P1–P4):

**P1 (Minimal baseline):**

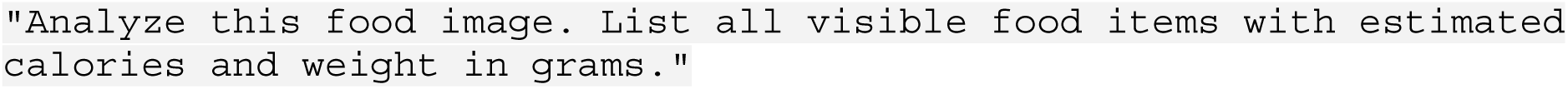

**P2 (Weight-first):** Instructs the model to estimate component weights in grams before calculating calories.

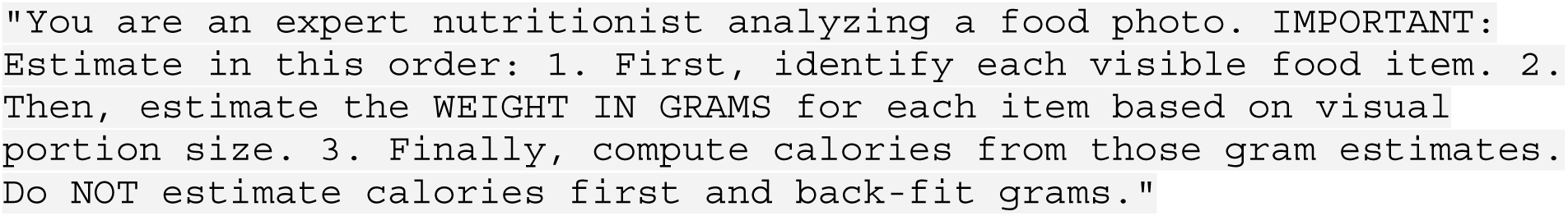

**P3 (Visible-portion):** Instructs the model to estimate weights and calories based on the actual visible portion in the photograph rather than defaulting to standard serving sizes.

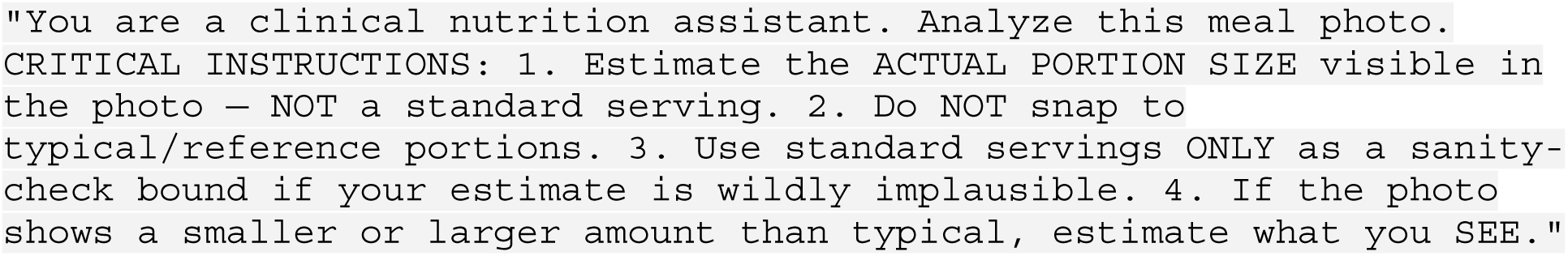

**P4 (Plausibility constraints):** Provides explicit plausibility bounds to help the model avoid unrealistic predictions.

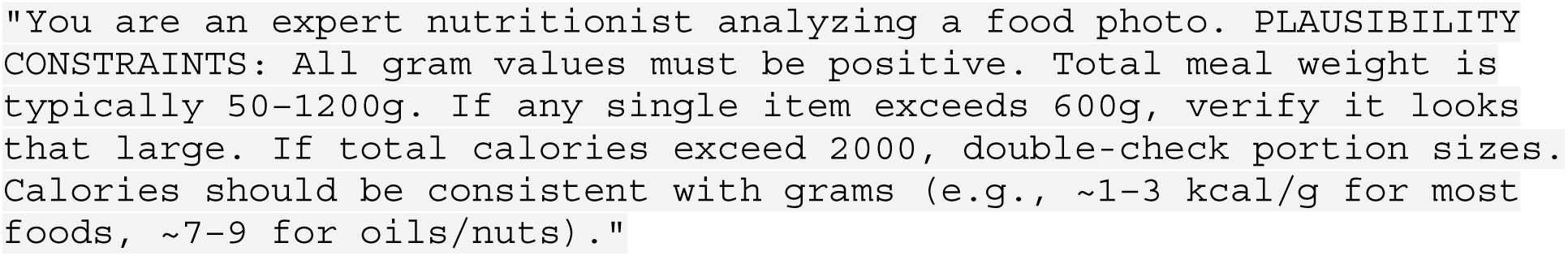

After identifying P3 as the best-performing single-pass variant, we defined a two-pass variant to test whether self-verification could improve upon it:

**P5 (Two-pass):** Uses P3 for the initial inference, then sends the model’s output back alongside the original image with a second prompt asking it to critique and refine its prediction:

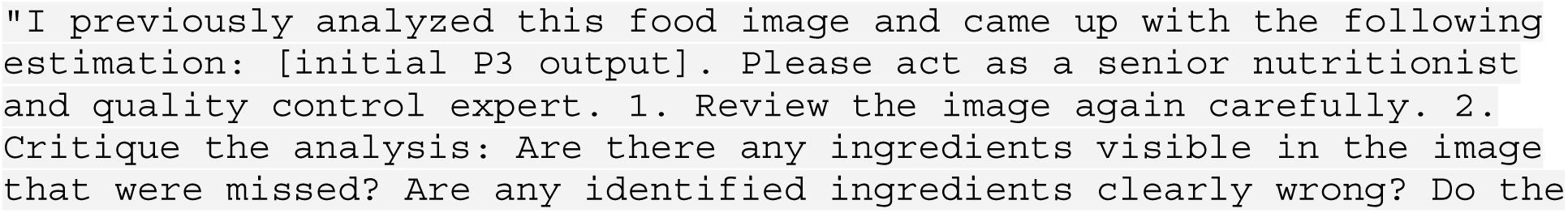
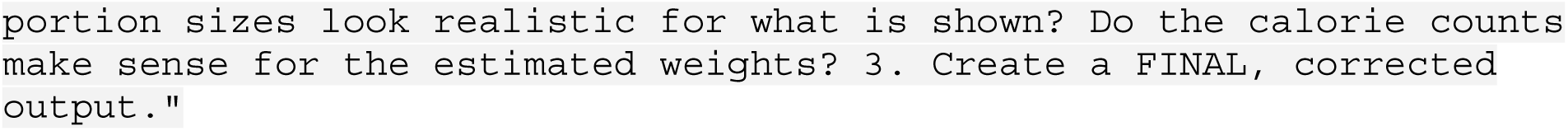

The best-performing single-pass prompt (P3) was then applied to all ten models for the full model comparison. We note that role descriptions varied across prompt variants (e.g., “expert nutritionist” in P2 and P4 vs. “clinical nutrition assistant” in P3); our ablation tested prompt strategies as complete packages rather than isolating individual factors such as role phrasing.

### Evaluation Metrics

To quantify model performance across our three tasks, calorie prediction, food weight estimation, and visible component recognition, we report the following set of metrics that capture both agreement with the original Nutrition5k labels and error magnitude. For calorie and weight estimation, we compute dish-level totals by summing predicted calories and predicted weights across all returned items and compare these totals to Nutrition5k’s dish-level labeled calories and total mass. For component recognition, we compare the predicted and original visible component sets using Jaccard similarity, described below.

- Concordance Correlation Coefficient (CCC): Lin’s CCC measures agreement between two continuous variables, accounting for both precision (correlation) and accuracy (systematic bias) [30]. Unlike Pearson’s r, CCC penalizes predictions that are correlated but systematically over- or under-estimated. Values range from −1 to 1, with 1 indicating perfect agreement. Since we report CCC as a descriptive agreement metric, significance testing is not conventionally applied.
- Mean Absolute Error (MAE): The average absolute difference between predicted and true values, expressed in kcal for calories and grams for weight.
- Mean Absolute Percentage Error (MAPE): MAE normalized by true values, expressed as a percentage. MAPE allows comparison of error magnitude across dishes of different sizes, but can be inflated by very small true values.
- Within-25% Accuracy: The percentage of predictions within 25% of the true value. This metric is commonly used in dietary assessment validation studies [31] and represents a practical benchmark for when predictions are “good enough” for dietary monitoring.
- Component Jaccard Index: For component detection, we compute Jaccard similarity at the token level between predicted and ground-truth visible component sets. Each ingredient phrase is lowercased, tokenized into individual words, and normalized by removing stopwords (e.g., “a,” “the,” “of”), cooking descriptors (e.g., “roasted,” “sliced”), color and size modifiers, and applying simple plural stemming (e.g., “tomatoes” → “tomato”) and synonym normalization (e.g., “aubergine” → “eggplant”). The resulting token sets from all predicted ingredients and all ground-truth ingredients are then compared using standard Jaccard similarity (|A ∩ B| / |A ∪ B|). We chose Jaccard over precision and recall because it provides a single measure that penalizes both missed components and hallucinated ones. We also removed a small set of clearly non-visual ingredients from the original labels (oils, vinegar, salt, spices, etc.) using a fixed stoplist; this affected 37 labeled components in total and did not change our reported metrics. It is important to note that this metric, like any image-based assessment, is fundamentally limited to visually identifiable components. Ingredients that are not visible in the image — such as added oils, vinegars, butter, dressings, and seasonings — cannot be reliably assessed through visual inspection alone, whether by a VLM or a human annotator. This is a well-recognized limitation of all image-based dietary assessment methods.

### Human Validation Study

To investigate the reliability of the Jaccard metric on Nutrition5k, we conducted a human validation study targeting our best-performing model and prompt combination (Gemini 2.0 Flash, P3). We developed a web-based annotation tool that presents annotators with food images alongside a merged ingredient list derived from both the original labels and AI labels. The tool operates in a source-hidden mode: for each ingredient, annotators see the item name and must classify it as approved (visibly present in the image), rejected (not present), or unsure (uncertain). Crucially, the annotator does not know which source predicted each ingredient, preventing bias toward either source.

Of the 3,229 images in our filtered dataset, 911 (28.2%) had identical original and AI food component sets; these images were excluded from manual review. From the remaining 2,318 images, a random subset was selected for manual review and divided among four annotators, each of whom independently reviewed a non-overlapping set of images. The annotation task required visual identification of food components (e.g., whether broccoli was visible in an image) rather than nutritional analysis. Annotators were not nutrition experts; the primary goal was to identify systematic errors in the Nutrition5k ground-truth labels rather than to produce expert-level annotations. Annotators had no prior exposure to the ground truth label–prediction pairings, and the annotation tool presented ingredients in a blinded, source-hidden format without indicating whether each item originated from the original Nutrition5k labels or from AI predictions, preventing bias toward either source. Annotator consistency was assessed by comparing approval rates across all four annotators, as reported in the Results. In total, 440 images (19% of eligible images) were reviewed, with each image assessed by a single annotator. This sample allowed us to examine systematic patterns in label quality across the dataset.

To merge original labels with the AI labels, we applied the same matching logic used in our primary evaluation pipeline. Items that matched between the two sources were presented once; unmatched items from either source were presented individually. After annotation, each ingredient’s approval status was linked back to its original source to enable source-specific computation of precision, recall, and Jaccard against the human-validated truth set.

### AI-Assisted Equivalency Matching

Our primary evaluation pipeline matches predicted and original label ingredients using token-overlap matching after stopword removal and lemmatization (i.e., reducing words to their base form, such as “tomatoes” to “tomato,” and removing common non-informative words such as articles and cooking descriptors). While effective for most cases, this approach fails for semantically equivalent foods with no shared tokens (e.g., “beef” vs. “steak,” “fish” vs. “salmon,” “yam” vs. “sweet potato”). To quantify the impact of these naming mismatches, we used GPT-4o- mini (OpenAI), selected for its low cost and strong performance on text classification tasks, to classify unmatched ingredient pairs as equivalent or not equivalent.

From the full 3,229-image dataset, we extracted all original label–AI ingredient pairs that (1) were not matched by our token-overlap algorithm, and (2) had zero token overlap under the pipeline’s matching logic (i.e., pairs that the original Jaccard computation could not have caught). This yielded 3,242 unique pairs. We used GPT-4o-mini to flag unmatched ingredient name pairs that refer to the same visible component, allowing synonyms and specificity differences (for example, ‘fish’ vs ‘salmon’) while rejecting merely similar or co-occurring foods. Pairs were processed in batches of 30 at temperature 0.0. Of 3,242 candidate pairs, 73 (2.3%) were classified as equivalent. We did not independently validate the accuracy of GPT-4o-mini’s equivalency classifications; misclassifications may introduce a small degree of error in the adjusted Jaccard scores.

### Ethical Considerations

This study analyzed publicly available food image data from the Nutrition5k dataset and did not involve human participants as research subjects. The annotation task involved visual inspection of food photographs by four annotators and did not collect personal or health-related data. No IRB approval was required.

## Results

### Prompt Engineering Study

Table 1 presents the prompt engineering results using Gemini 2.0 Flash across all 3,229 images. P3 achieved the best overall performance, with calorie CCC of 0.744 and 42.0% of predictions within 25% of the original labels. The improvement from our baseline P1 prompt to P3 increased calorie CCC from 0.693 to 0.744 (an absolute gain of 0.051, or 7.4% relative) and improved within- 25% accuracy from 30.8% to 42.0% (a 36% relative improvement) (Fig. 2). This is potentially because VLMs suffer from portion-size bias, and snap to canonical serving sizes (i.e. 1 cup of rice) regardless of what is actually on the plate.

**Fig. 1.**
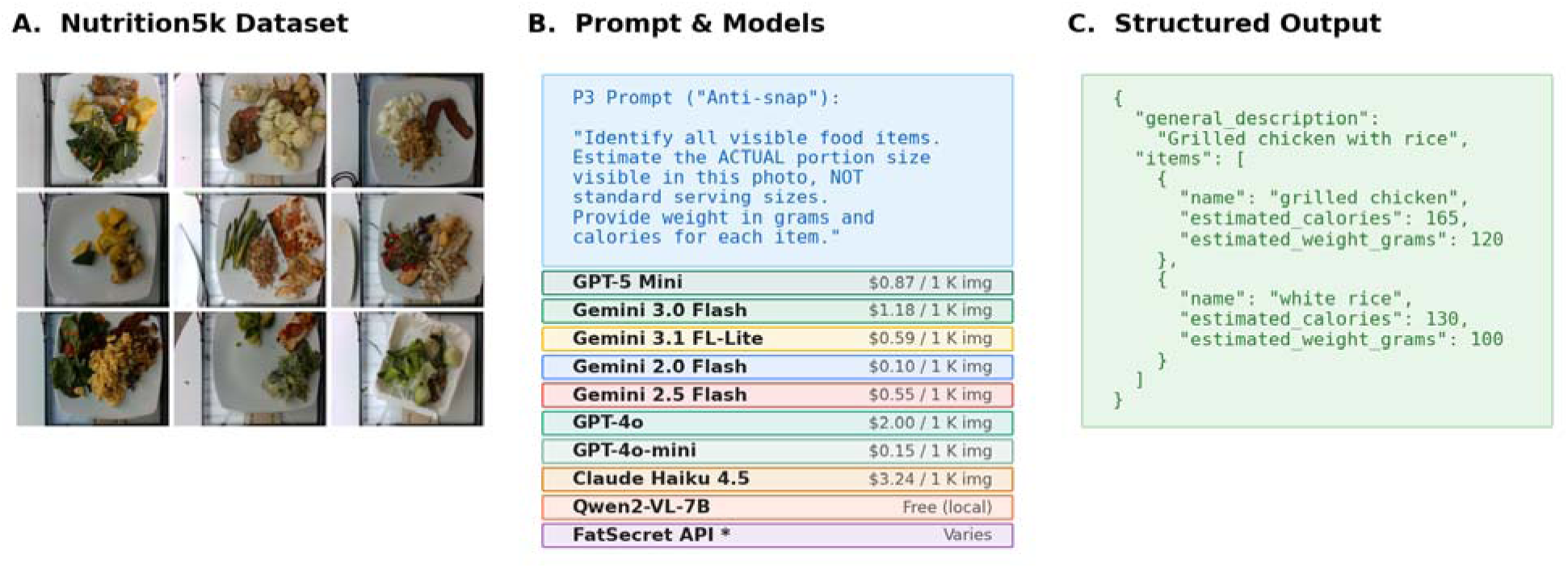
**(A)** Representative food images from the Nutrition5k dataset. **(B)** The standardized P3 prompt and the ten models evaluated, with estimated cost per 1,000 images. **(C)** Example structured JSON output showing identified components, estimated weight, and predicted calories. *FatSecret API uses a fixed recognition pipeline with no customizable prompt input.

**Fig. 2.**
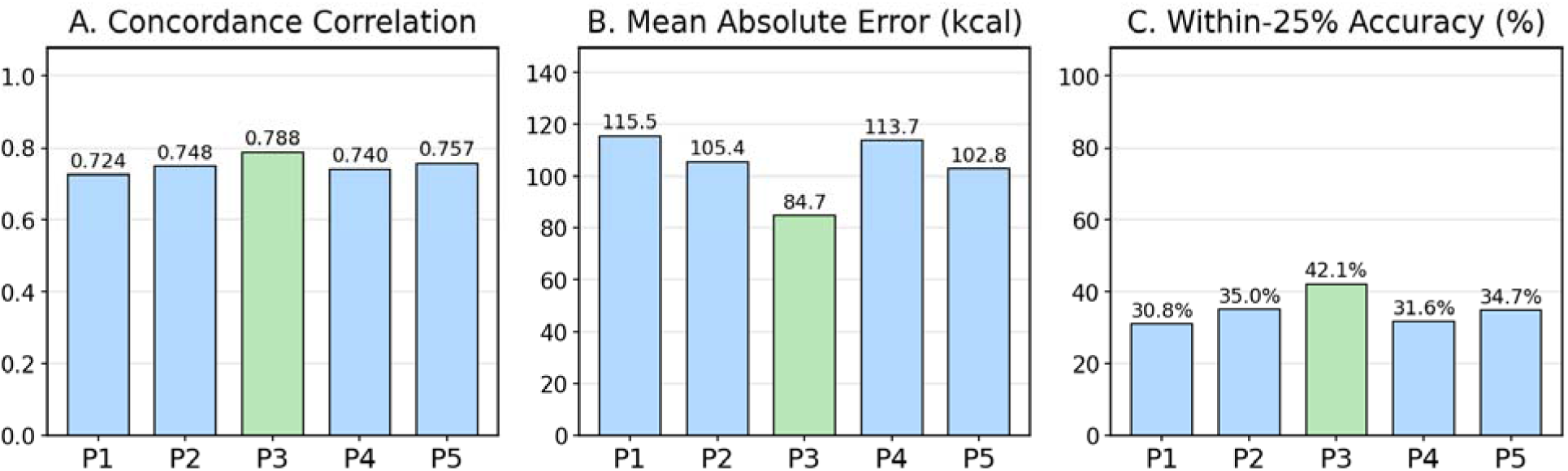
Bar chart with prompt engineering study results showing (**A**) Calorie CCC, **(B)** Calorie MAE, and (**C)** Within-25% accuracy across five prompt variants.

**Table 1:** Prompt engineering study results (Gemini 2.0 Flash, N=3,229). Bold values indicate best performance in each column.

| Prompt | Description | Cal CCC | Cal MAE | Within 25% | Wt CCC | Wt MAE | Jaccard |
| --- | --- | --- | --- | --- | --- | --- | --- |
| **P3** | **Visible portions** | **0.744** | **85.9** | **42.0%** | **0.750** | **72.0** | **0.629** |
| P5 | Two-pass | 0.723 | 103.9 | 34.7% | 0.710 | 86.9 | 0.624 |
| P2 | Weight-first decomposition | 0.715 | 106.5 | 34.9% | 0.687 | 94.3 | 0.627 |
| P4 | Explicit constraints | 0.710 | 114.9 | 31.6% | 0.702 | 91.6 | 0.621 |
| P1 | Minimal baseline | 0.693 | 117.0 | 30.8% | 0.689 | 91.9 | 0.622 |

Conversely, while two-pass prompting approaches have shown promise in other domains for reducing errors through self-correction, [32–33] our two-pass prompting approach (P5) yielded results that were equal to or worse than single-pass inference across all metrics. This is potentially because two-pass self-correction helps most on problems with checkable constraints or external feedback, but calorie/weight estimation from a single image is largely non-verifiable, so a second pass often adds noise or pulls predictions toward generic “typical” weights.

### Model Comparison on Full Dataset

For calorie estimation, Gemini 3.0 Flash achieved the highest CCC (0.767), followed by GPT-5 Mini (0.760) and Gemini 3.1 Flash Lite (0.754) (Table 2) (Suppl. Fig. 1). For weight estimation, Gemini 3.0 Flash achieved the highest weight CCC (0.779), followed by Gemini 3.1 Flash Lite and GPT-5 Mini (both 0.772) (Suppl. Fig. 2). All VLM models demonstrated reasonable component detection capabilities, with mean Jaccard scores ranging from 0.386 (Claude Haiku 4.5) to 0.655 (Gemini 3.1 Flash Lite) among VLMs (Fig. 3).

**Fig. 3.**
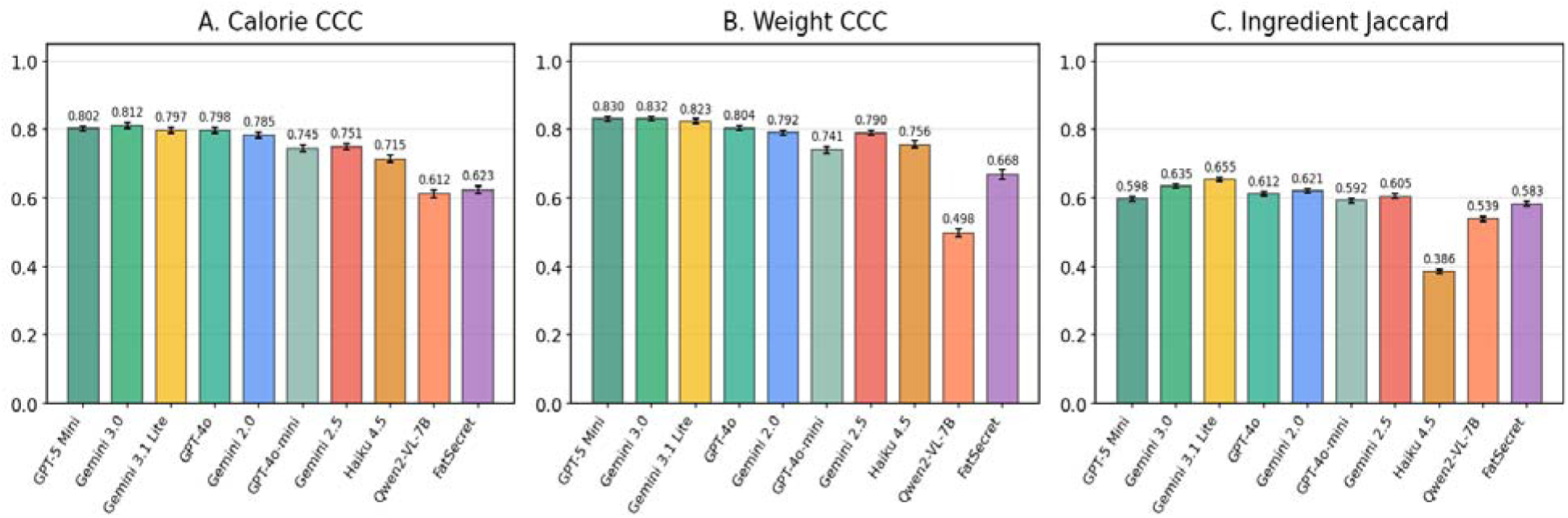
Bar chart comparing model performance across key metrics: **(A)** Calorie CCC, **(B)** Weight CCC, and **(C**) Mean component Jaccard.

**Table 2:** Performance comparison on full Nutrition5k filtered dataset (N=3,229) using P3 prompt. Best values are highlighted in bold. Note that FatSecret only processed 3,216 of our 3,229 images due to an API error on 13 images. Performance for Gemini 2.0 Flash with model P3 may differ slightly between Table 1 and Table 2 as these results were generated from a separate inference run. †GPT-5 Mini processed 3,016 of 3,229 images; 213 returned unparseable responses. Cost/1K denotes the estimated cost (USD) to process 1,000 images using published API pricing at the time of evaluation (January 2026); pricing may change. [14,34] FatSecret pricing is plan and usage dependent, so we report it as Varies. *Qwen2-VL-7B is open-source and free to run, but requires infrastructure/compute costs.

| Model | N | Cal CCC | Cal MAE | Cal MAPE | Within 25% | Wt CCC | Wt MAE | Jaccard | Cost/1K |
| --- | --- | --- | --- | --- | --- | --- | --- | --- | --- |
| Gemini 3.0 Flash | 3,229 | **0.767** | **80.7** | 44.4% | 43.7% | **0.779** | 53.8 | 0.635 | \$1.18 |
| GPT-5 Mini | 3,016 | 0.760 | 87.8 | 52.0% | 39.8% | 0.772 | **53.7** | 0.598 | \$0.87 |
| Gemini 3.1 Flash Lite | 3,229 | 0.754 | 82.5 | **39.8%** | **44.5%** | 0.772 | 55.1 | **0.655** | \$0.59 |
| GPT-4o | 3,229 | 0.752 | 82.9 | 41.7% | 42.1% | 0.752 | 60.4 | 0.612 | \$2.00 |
| Gemini 2.0 Flash | 3,229 | 0.742 | 87.9 | 46.5% | 41.3% | 0.752 | 73.7 | 0.621 | \$0.10 |
| Gemini 2.5 Flash | 3,229 | 0.718 | 111.1 | 60.7% | 35.9% | 0.749 | 72.4 | 0.605 | \$0.55 |
| GPT-4o-mini | 3,229 | 0.704 | 94.6 | 53.8% | 36.5% | 0.692 | 67.1 | 0.592 | \$0.15 |
| Claude Haiku 4.5 | 3,229 | 0.680 | 105.5 | 67.3% | 34.8% | 0.716 | 73.0 | 0.386 | \$3.24 |
| Qwen2-VL-7B | 3,229 | 0.612 | 121.0 | 120.4% | 30.1% | 0.497 | 97.6 | 0.539 | Free* |
| FatSecret | 3,216 | 0.597 | 149.6 | 87.9% | 28.5% | 0.612 | 92.3 | 0.583 | Varies |

**Table 3:** Comparison of recent VLM-based nutrition studies on Nutrition5k. We report calorie mean absolute error (MAE) and mean absolute percentage error (MAPE) when available, and ingredient-set overlap (Jaccard) when reported. Ingredient-overlap metrics are not directly comparable across studies due to differences in label mappings and evaluation procedures.

| Study | Calories MAE (kcal) | Calories MAPE (%) | Component metric (ingredient overlap) |
| --- | --- | --- | --- |
| Our study | 80.7 (Gemini 3.0 Flash) | 44.4 (Gemini 3.0 Flash) | Jaccard: ~0.82 (Gemini 2.0 with P3, with extrapolation) |
| **Tanabe & Yanai** | 82.7 (GPT-4o)<br>78.8 (GPT-4o + volume) | 46.7 (GPT-4o)<br>43.4 (GPT-4o + volume) | Not reported |
| **Khlaisamniang et al.** | 88.86 (standard)<br>80.32 (two-step) | Not reported | **Jaccard** (USDA / Nutrition5k mapping): 0.312 / 0.330 (standard) - 0.338 / 0.338 (two-step) |

### Fruit and Vegetable Recognition

Given the clinical importance of fruit and vegetable intake monitoring, we conducted a subanalysis of model performance on dishes containing these components. Of our 3,229 images, 2,031 (63%) contained at least one fruit or vegetable. Gemini 2.0 Flash (P3) achieved significantly higher Jaccard scores on fruit- and vegetable-containing dishes (mean 0.655) compared to non-fruit/vegetable dishes (mean 0.560; P<.001, Welch’s t-test). When isolating only fruit and vegetable components, the model achieved a mean Jaccard of 0.730 for fruits (median 1.0, indicating perfect recognition on most dishes) and 0.629 for vegetables. These scores are computed against the original Nutrition5k labels; as demonstrated in the validation study below, actual model accuracy is likely higher due to systematic omissions in the ground truth annotations.

### Food component recognition validation

To investigate the relatively modest Jaccard scores observed across all models (0.54–0.62), we manually inspected a subset of images with low component recognition (Fig. 4). We identified two distinct categories of error: true model failures, where the AI genuinely misidentified or missed food items, and false failures, where the AI was penalized despite producing correct or reasonable predictions.

**Fig. 4.**
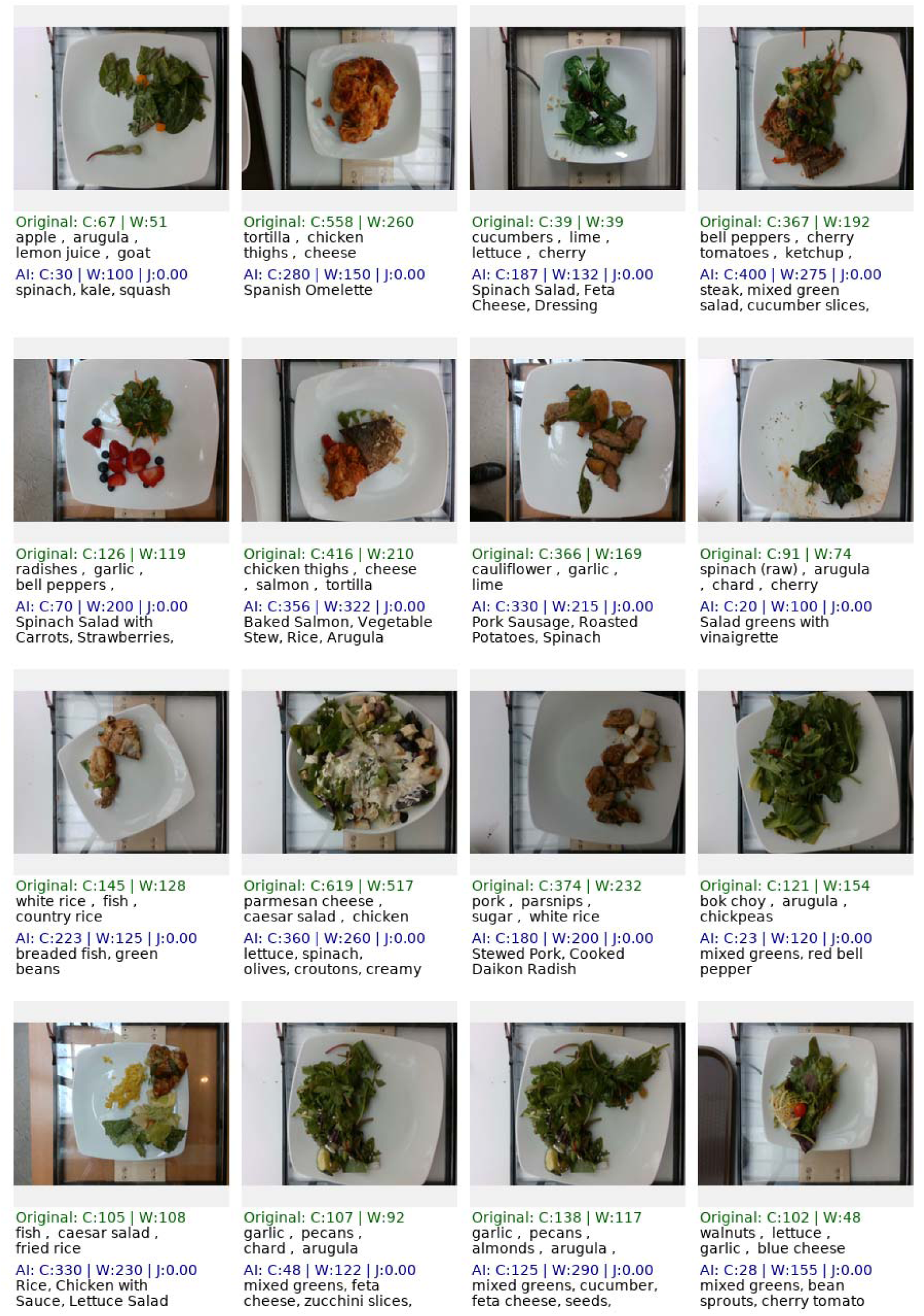
Montage of dishes with low component recognition. Each panel shows the food image with original labels and AI-predicted components, along with calorie **(C)** and weight **(W**) estimates, and Jaccard score **(J).**

**True failures:** These reflect genuine limitations of the model:

1. Component confusion: Visually similar foods are frequently misidentified (e.g., yam vs. sweet potato vs. butternut squash; different types of leafy greens).
2. Partial visibility: Components obscured by sauces, overlapping items, or presentation angle are often missed or misidentified.

**False failures:** These reflect limitations of the evaluation rather than the model:

1. Naming mismatches: Semantically equivalent ingredients with no shared tokens (e.g., “beef” vs. “steak,” “fish” vs. “salmon”) are treated as non-matches by the token-overlap metric, artificially deflating Jaccard scores.
2. Annotation errors: For more complex dishes, certain visually identifiable components were omitted from the original labels, or in some rare cases, components not present in the image were included.

To address the false failures, we conducted two analyses. First, we prompted GPT-4o-mini (temperature 0.0) to classify whether two food component names refer to the same item, answering Y or N for each pair. The prompt accepted synonyms (“cilantro”/“coriander”), spelling variants (“brussel sprouts”/“brussels sprouts”), different forms of the same item (“beef”/“steak”), and parent-child relationships (“fish”/“salmon”), but explicitly rejected visually similar but distinct foods, foods in the same category, foods containing another food as an ingredient, and foods commonly served together.

Applying the 73 AI-discovered equivalency pairs to the full dataset increased the mean Jaccard for Gemini 2.0 Flash from 0.621 to 0.635. Of 3,242 candidate pairs with zero token overlap, 73 (2.3%) were classified as equivalent (Fig. 5). This modest improvement indicates that naming mismatches account for only a small fraction of the gap between reported and true model performance.

**Fig. 5.**
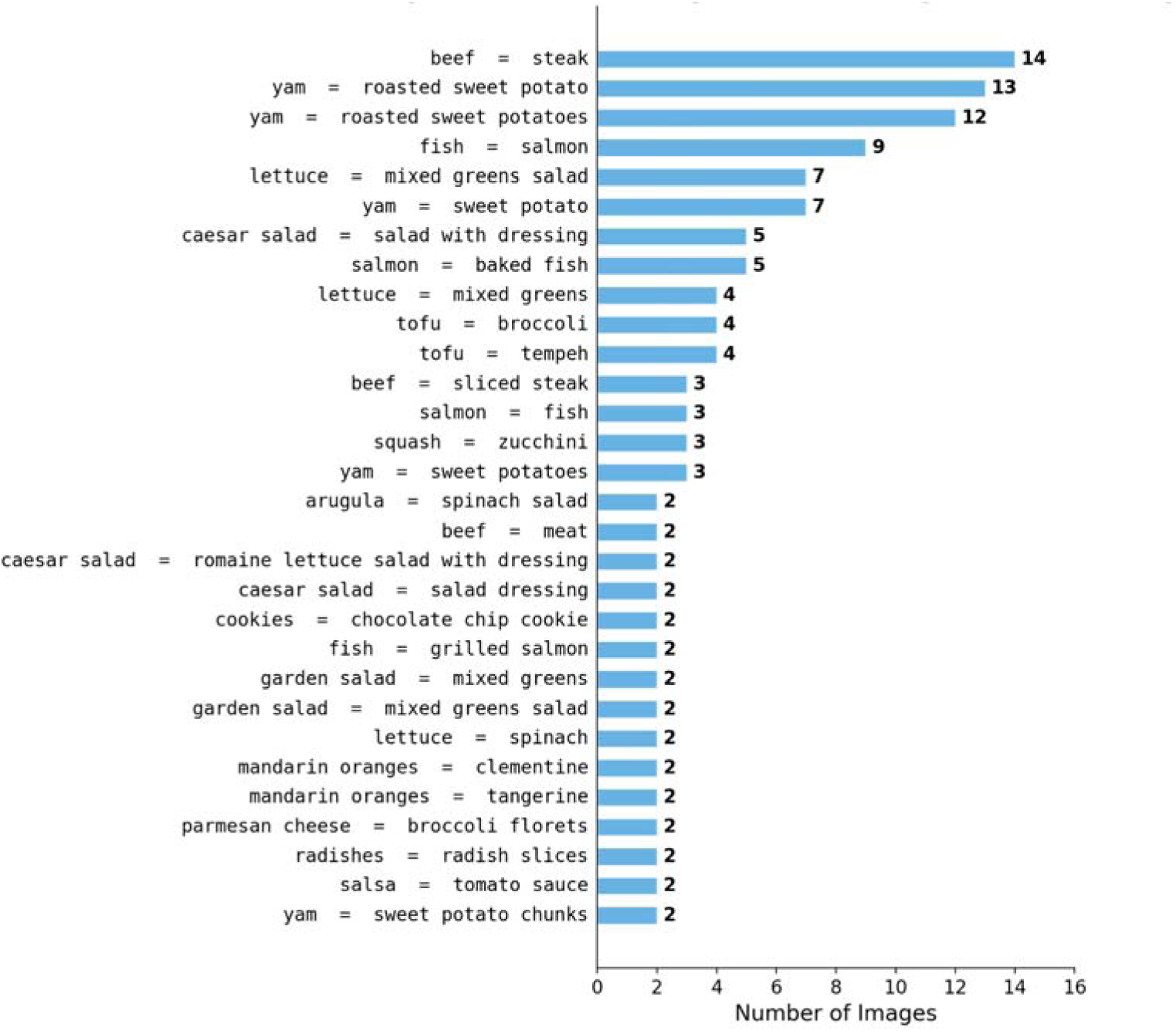
Bar chart showing top 30 food component equivalencies between the original and AI-predicted set, as identified by GPT-4o-mini.

Second, we developed a web-based annotation tool in which four human annotators reviewed food images alongside a merged list of components from both the original Nutrition5k labels and AI predictions (Gemini 2.0 Flash, prompt P3). For each component, annotators indicated whether it was visibly present in the image (Fig. 6). Annotators were not able to see whether a particular food component came from the original labels or from the AI labels. Annotators might mark a food component as correct (visibly present), incorrect (not present), or unsure. Four annotators each reviewed a non-overlapping subset of images, for a total of 440 images. 911 images already had matching labels and were thus excluded from review.

**Fig. 6.**
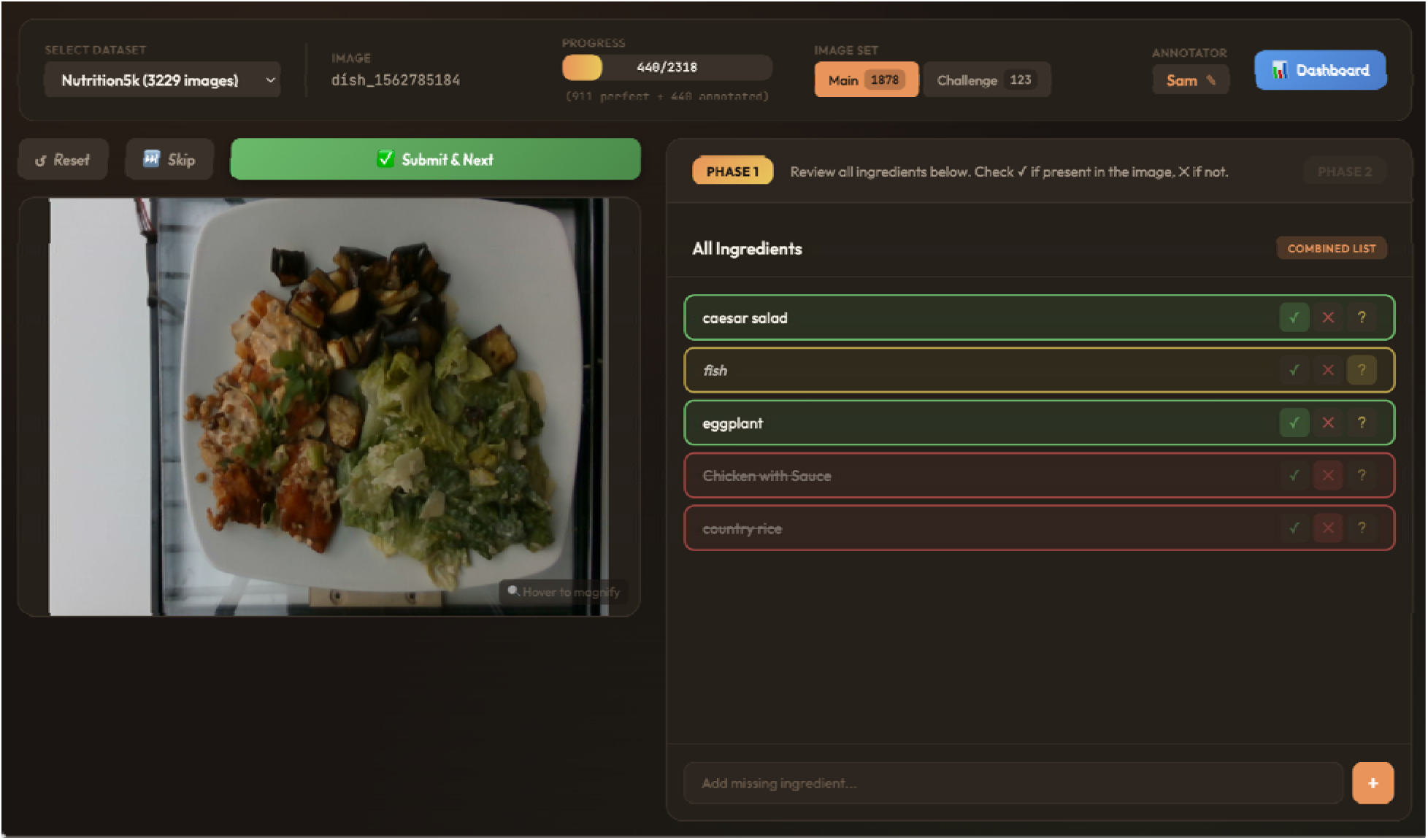
Screenshot showing our web-based annotation tool, which allows annotators to determine the accuracy of both the original and AI-predicted food component labels.

From the list of original ingredients, annotators approved 77.9% (804 of 1,032), rejected 12.5% (129), and marked 9.6% (99) as unsure. From the list of AI-predicted ingredients, annotators approved 84.5% of items (1,244 of 1,473), rejected 9.4% (138), and marked 6.2% (91) as unsure (Fig. 7c). Notably, among ingredients that the original labels did not include but the AI did (n=453), annotators approved 56.3% (255) as present, indicating that the AI identified a substantial number of real food items that the original annotators missed. Under the strict interpretation (unsure = incorrect), the AI achieved precision of 83.8%, recall of 82.7%, and Jaccard of 72.6%, compared to 80.2%, 54.7%, and 48.0% for the original labels. The AI’s substantially higher recall (82.7% vs. 54.7%) reflects the Nutrition5k’s original annotators’ tendency to omit visible food items (Fig. 7a).

**Fig. 7.**
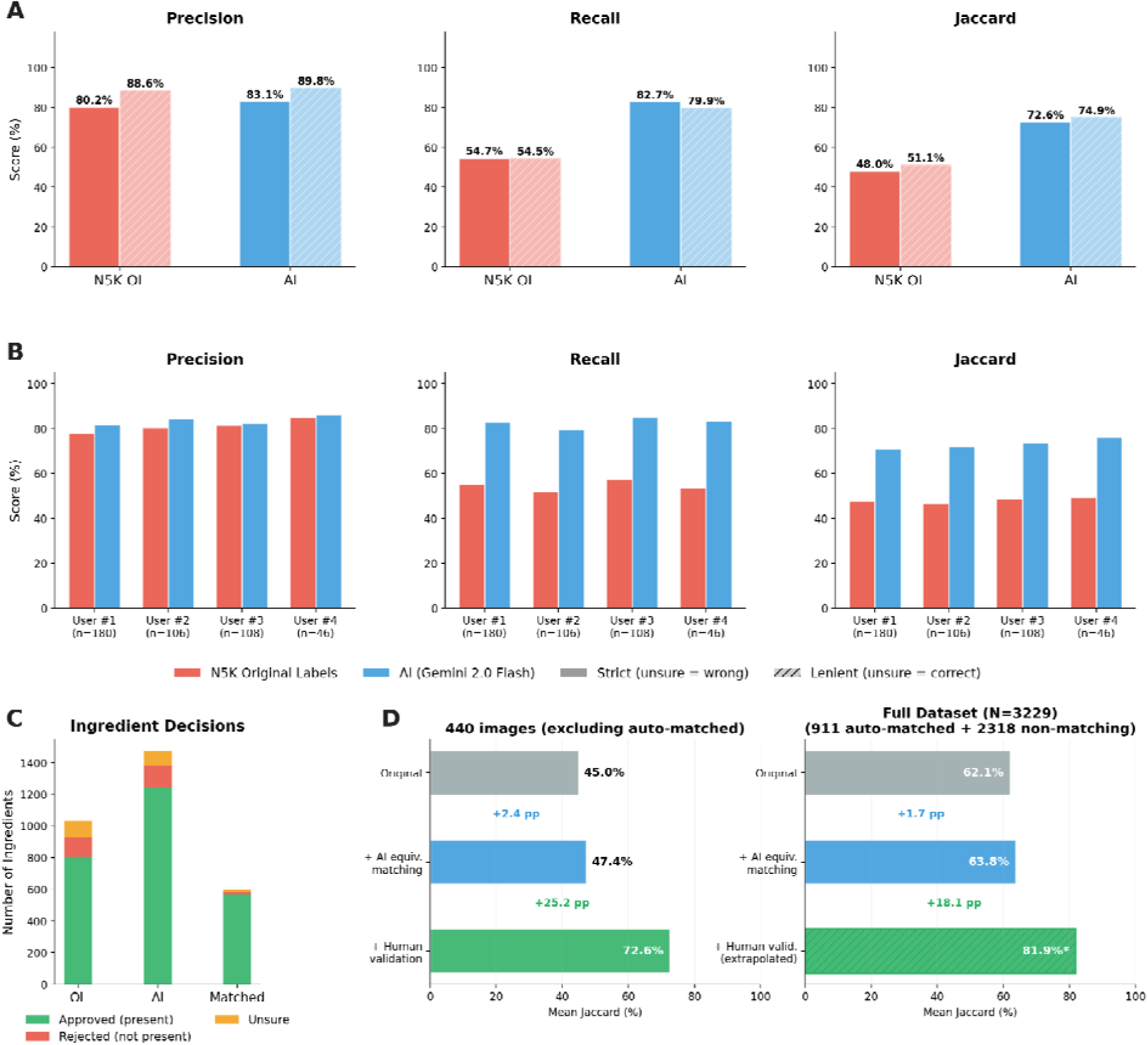
Human validation comparing N5k’s original labels with AI-predicted ingredients (A) Bar charts comparing precision, recall, and Jaccard. Hatched bars show lenient interpretation (unsure items treated as correct). (B) Per-annotator comparison of precision, recall, and Jaccard. (C) Annotator decisions based on whether the identified component was present. (D) Cumulative Jaccard improvement after AI equivalency matching (blue) and human label correction (green) for 440 validated images (left) and extrapolated to the full dataset (right). Auto-matched refers to images where the original and AI labels already match; these images were excluded from the web-based annotation tool.

On the 440 validated images, Gemini 2.0 Flash (using prompt P3) achieved a mean Jaccard of 0.726 against the human-validated labels, compared to 0.450 against the original labels on the same images. To extrapolate this correction to the full dataset, we note that 911 of 3,229 images (28.2%) already have perfect agreement (Jaccard = 1.0), and the reported full-dataset mean is 0.621 (Table 2), implying a mean of 0.472 across the remaining 2,318 non-perfect images. We computed two extrapolations: an additive correction (applying the observed delta of +0.276 to the non-perfect mean) and a multiplicative correction (scaling the non-perfect mean by the observed ratio 0.726/0.450 = 1.61). These yield estimated full-dataset Jaccard scores of 0.82 and 0.84, respectively (Fig. 7d). We report the more conservative additive estimate of ∼0.82. Because the validated sample was drawn from images with discrepant labels, which may represent harder cases, this figure should be interpreted as an approximate upper-bound estimate rather than a precise correction. Our validation study was conducted using Gemini 2.0 Flash as it was the most cost-effective model at the time of study. Given that Gemini 3.1 Flash Lite achieves a higher raw Jaccard (0.655 vs. 0.621), we expect its validated Jaccard to be even higher than the 0.82 reported here.

To assess whether our findings reflected individual annotator bias, we compared approval rates across all four annotators (Fig. 7b). Each annotator reviewed a non-overlapping subset of images, so differences in aggregate metrics can reflect variation in image difficulty as well as annotator stringency. Nonetheless, approval rates were similar across annotators for both the original ingredients and the AI-predicted ingredients, and all four annotators independently showed higher approval rates for the AI-derived labels compared with the original Nutrition5k labels.

Because each annotator reviewed a non-overlapping image set, standard inter-annotator agreement statistics (e.g., Fleiss’ kappa) could not be computed; the consistency of per-annotator approval rates serves as a qualitative check rather than a formal reliability measure.

### Comparison with Prior Work

In the original Nutrition5k study, the supervised baseline model trained on only the single overhead RGB image achieved a calorie MAE of 70.6 kcal. Our best performing VLM (Gemini 3.0 Flash, prompt P3) uses only the overhead RGB image and no Nutrition5k training and achieves a competitive calorie MAE of 80.7 kcal. The Nutrition5k study showed that including the matching overhead depth map, reduces error to 47.6 kcal, and using a depth-derived volume estimate (“volume scalar”) reduces it further to 41.3 kcal. This demonstrates that weight estimation is substantially easier when 3D information is available. The researchers did not evaluate food component detection accuracy. [24]

Using Nutrition5k, Tanabe and Yanai benchmarked GPT-4o and reported a MAE of 82.7 kcal (MAPE 46.7%). When they added a “volume injection” module (object detection/segmentation plus depth estimation), GPT-4o improved to 78.8 kcal MAE (MAPE 43.4%). [16] Similarly, Khlaisamniang et al. tested a two-step, “nutritionist-style” prompting workflow on Nutrition5k and saw GPT-4o improve from 88.86 to 80.32 kcal MAE, with small gains in their component Jaccard scoring. [35]

## Discussion

Our evaluation demonstrates that current VLMs can perform automated dietary assessment with reasonable accuracy. Gemini 3.0 Flash achieved the best calorie estimation (CCC=0.767, MAE=80.7 kcal), while Gemini 3.1 Flash Lite offered very comparable performance (CCC=0.754, MAE=82.5 kcal) with the highest ingredient recognition (Jaccard=0.655) at roughly half the cost ($0.59 vs. $1.18 per 1,000 images). Among earlier-generation models, Gemini 2.0 Flash remained competitive (CCC=0.742) at just $0.10 per 1,000 images, though both Gemini 2.0 Flash and 2.5 Flash have been marked for deprecation by Google, and model availability should be verified at time of deployment. GPT-4o showed strong performance (CCC=0.752) but at considerably higher cost ($2.00/1,000 images). Our prompt engineering study revealed that explicit instructions to estimate “the actual visible portion” (P3) improved calorie CCC by 7.4% over baseline. This finding suggests that VLMs, when not explicitly instructed otherwise, default to canonical serving sizes rather than estimating the actual visible portion, a behavior analogous to the portion-size anchoring seen in traditional dietary assessment methods. These findings also suggest that researchers and developers deploying VLMs for dietary assessment should invest in prompt optimization, with prompts co-developed in collaboration with nutrition researchers and registered dietitians. Nonetheless, calorie and weight estimation errors remained high across all models (MAPE 39.8–120.4%; weight MAE 53.7–97.6g), even when the correct food components were identified. Additionally, GPT-5 Mini failed to return parseable responses for 213 of 3,229 images (6.6%), suggesting that practical deployment reliability varies across providers and should be evaluated alongside accuracy.

Recent VLM nutrition papers mostly agree on the general findings: models usually identify the obvious foods correctly, but they struggle when the job turns into portion sizing, mixed dishes, or anything “hidden” like oil, sauces, or fillings. In a controlled photo study with small, medium, and large portions, Fridolfsson et al. found that GPT-4o and Claude 3.5 Sonnet had similar errors (about mid-30% MAPE for both weight and energy), and all models increasingly underestimated calories and weight as portions got larger. [15] A study by Cinar et al. that focused only on GPT-4o reported the same challenge: accuracy was worse on complex meals and only improved after the researchers provided extra context about components. [36] Bigger food recognition and captioning benchmarks reinforce the same pattern we see in our analyses: Food-500Cap shows that fine-grained food descriptions remain hard and performance varies by cuisine/region. [37] These limitations are also common in traditional dietary assessment methods such as food records and food frequency questionnaires, highlighting the need for more research and development in this area.

While all models achieved component detection Jaccard scores between 0.39 and 0.66, our validation study reveals that these scores substantially underestimate true model performance. We find that the reported Jaccard scores are depressed by two factors: naming mismatches in the evaluation metric, and systematic quality issues in the Nutrition5k annotations. After correction, we extrapolate the corrected Jaccard for Gemini 2.0 Flash to approximately 0.82, though this estimate assumes the 440 validated images are representative of the broader non- matching set. Because Gemini 3.1 Flash Lite achieves a higher raw Jaccard than 2.0 Flash (0.655 vs. 0.621), its true validated score may exceed this estimate. These findings highlight a broader challenge in food image benchmarking: constructing reliable labels for complex, multi- component meals is inherently difficult. Lastly, it’s important to note that Nutrition5k primarily consists of Western-style cafeteria meals with relatively clear presentation, taken using overhead cameras. It may be beneficial to perform further study on home-cooked meals, restaurant dishes, or culturally diverse cuisines where components and portion sizes are not always clearly visible from overhead images.

While our evaluation focused on VLM performance for food recognition, weight estimation, and calorie prediction, commercially available AI-powered nutrition apps such as Keenoa [12] and Rx Food [13] already provide richer nutritional outputs, including macronutrient and micronutrient breakdowns, by linking recognized foods to curated nutrient databases. The VLM outputs evaluated here are limited to ingredient identification, weight, and calorie estimates. However, VLM-generated food identifications could in principle be paired with standard nutrient databases to derive more comprehensive nutritional profiles, though this is beyond the scope of the present study. Additionally, image-based dietary assessment — whether by VLMs, commercial apps, or human annotators — is inherently limited to visually identifiable components. Hidden ingredients such as added oils, butter, dressings, and seasonings cannot be detected from a photograph alone, representing a fundamental constraint shared across all image-based approaches. Our evaluation also did not include beverages, which present their own challenges for image-based assessment, including distinguishing diet from regular drinks and identifying additions to coffee and tea. Our prompt engineering study was conducted using Gemini 2.0 Flash only; the optimal prompt may differ across model families, and applying P3 uniformly may understate the potential of non-Gemini models. Finally, deploying VLM-based dietary assessment in practice requires transmitting food photographs to third-party API providers, which raises privacy considerations that must be addressed before clinical use.

In summary, we demonstrate that off-the-shelf VLMs can perform automated dietary assessment from single overhead photographs with accuracy approaching supervised baselines, at low per-image cost. Among the models tested, Gemini 3.0 Flash offers the best overall accuracy while Gemini 3.1 Flash Lite provides the best cost-performance tradeoff, achieving near-equivalent accuracy with the highest ingredient recognition at the lowest per-image cost among top performers. Our validation study, conducted using Gemini 2.0 Flash (the most cost-effective model at the time of study), demonstrates that true ingredient recognition far exceeds what raw Jaccard scores suggest. Targeted prompt engineering yields meaningful accuracy gains across all models. These findings support the feasibility of deploying VLMs in scalable dietary monitoring applications; however, further model refinement is needed to improve accuracy. Our study shows that VLMs have promise in tasks such as identifying foods and ingredients and generating reasonably accurate estimates of food weight and energy content. Still, the ultimate value of these tools for nutrition research will depend on whether these image-based outputs can be

translated into meaningful measures of nutrient intake and diet quality. If successful, automated dietary assessment will have great potential for clinical and research applications in a broad range of health conditions. Realizing this potential will also depend on whether these approaches prove feasible to implement at scale in clinical practice and population-based studies.

## Appendix

**Supplementary Figure 1.**
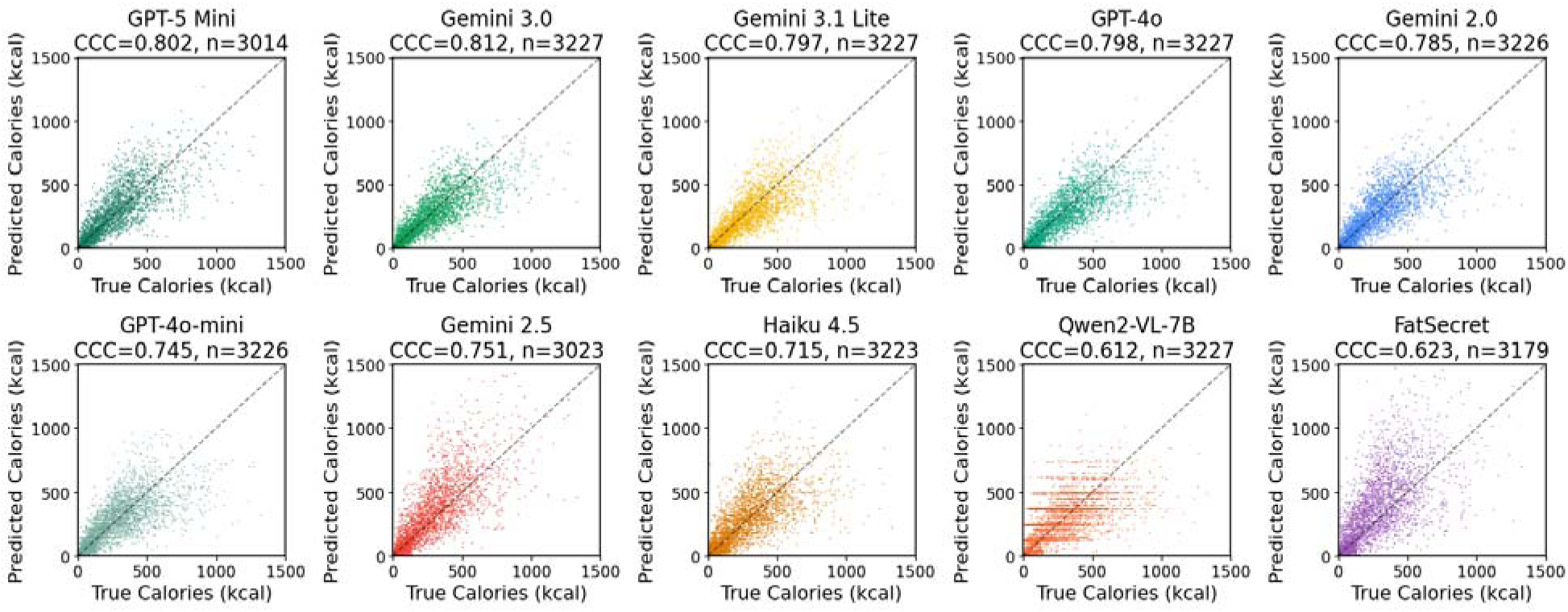
Scatter plots showing AI-predicted calories (y-axis) against original calories (x-axis) for all 3,229 images across our ten models.

**Supplementary Fig. 2.**
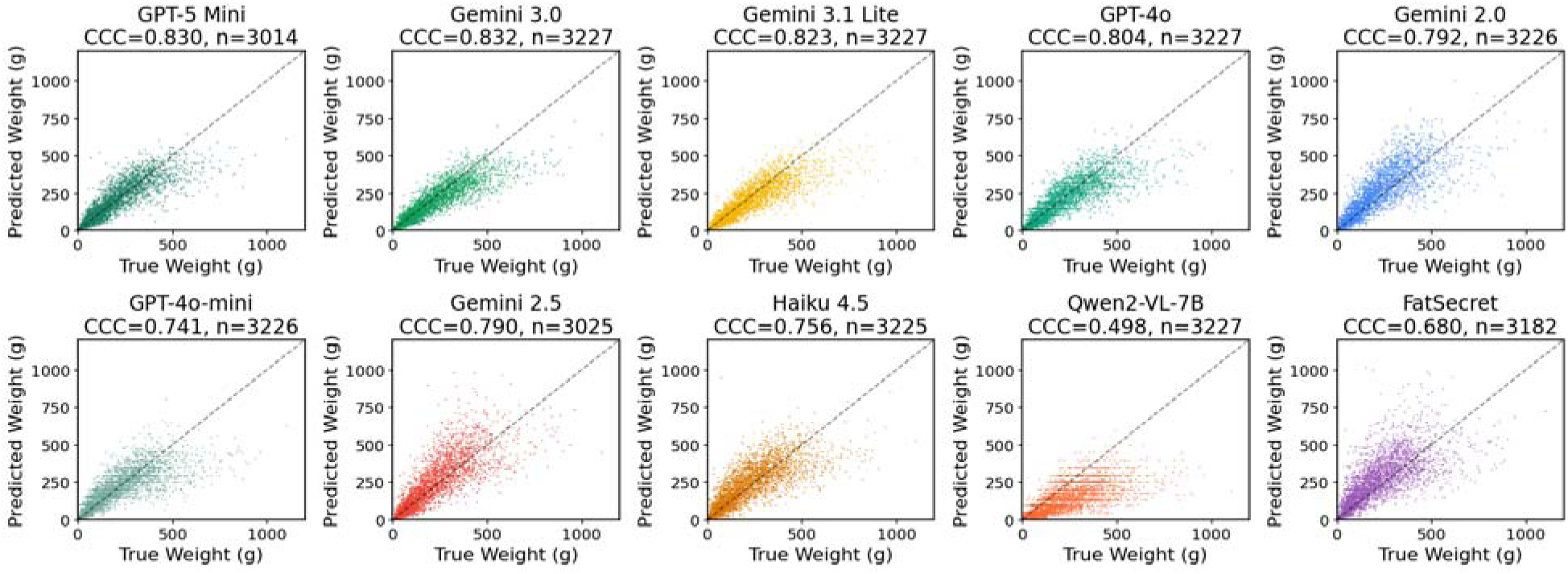
Scatter plots showing predicted weight (y-axis) against original weight (x- axis) for all 3,229 images across our ten models.

## Acknowledgments

Generative AI tools, including Claude (Anthropic), were used for copy editing, formatting, revising draft text, and assisting with the development of evaluation and analysis code. The authors have reviewed all AI-assisted output and take full responsibility for the content of this manuscript.

## Data Availability

The Nutrition5k dataset is publicly available at https://github.com/google-research-datasets/Nutrition5k.

## Code Availability

Our code is available on our Github at https://github.com/charterer/vlm-food-benchmark.

## Authors’ Contributions

SS designed the study, developed the evaluation pipeline, ran all experiments, analyzed the data, and wrote the manuscript. AT supervised the study and provided input on direction. All other authors contributed feedback and manuscript revisions.

## Acknowledgment

We thank Antonios Tsirigos and Anna Karampachtsi for systematically comparing ingredient lists with the contents depicted in the meal photos using a web-based annotation tool.

## Abbreviations

API: application programming interface
CCC: concordance correlation coefficient
GPU: graphics processing unit
HPC: high-performance computing
JSON: JavaScript Object Notation
MAE: mean absolute error
MAPE: mean absolute percentage error
RGB: red-green-blue
VLM: vision- language model

## References

1. Subar, A. F. et al. Using intake biomarkers to evaluate the extent of dietary misreporting in a large sample of adults: the OPEN study. Am. J. Epidemiol. 158, 1–13 (2003). https://pubmed.ncbi.nlm.nih.gov/14559779/

2. Poslusna, K., Ruprich, J., de Vries, J. H., Jakubikova, M. & van’t Veer, P. Misreporting of energy and micronutrient intake estimated by food records and 24 h recalls. Br. J. Nutr. 101(Suppl 2), S73–S85 (2009). doi:10.1017/S0007114509990602. https://pubmed.ncbi.nlm.nih.gov/19594967/

3. Boushey, C. J. et al. New mobile methods for dietary assessment: review of image-assisted and image-based dietary assessment methods. Proc. Nutr. Soc. 76, 283–294 (2017). 10.1017/s0029665116002913

4. Grand View Research. Diet and Nutrition Apps Market. (accessed 2026-01-29). https://www.grandviewresearch.com/press-release/global-diet-nutrition-apps-market

5. Bossard, L., Guillaumin, M. & Van Gool, L. Food-101: Mining discriminative components with random forests. In Proc. Eur. Conf. Comput. Vis. (ECCV) (2014) doi:10.1007/978-3-319-10599-4_29. https://link.springer.com/chapter/10.1007/978-3-319-10599-4_29

6. Min, W. et al. A survey on food computing. ACM Comput. Surv. 52, 1–36 (2019). arXiv:1808.07202. https://arxiv.org/abs/1808.07202

7. Myers, A. et al. Im2Calories: Towards an Automated Mobile Vision Food Diary. In Proc. IEEE Int. Conf. Comput. Vis. (ICCV) (2015). doi:10.1109/ICCV.2015.146. https://www.semanticscholar.org/paper/115f30924be576b00ad5184c9ba963430887ba4f

8. Fang, S., Shao, Z., Mao, R., Fu, C., Delp, E. J. & Zhu, F. Single-View Food Portion Estimation: Learning Image-to-Energy Mappings Using Generative Adversarial Networks. In Proc. 2018 25th IEEE Int. Conf. Image Process. (ICIP), 251–255 (2018). doi:10.1109/ICIP.2018.8451461. https://www.semanticscholar.org/paper/3ab1450b2db38e95eb1ba21bcd577b051cec4c56

9. Yang, Y., Jia, W., Bucher, T., Zhang, H. & Sun, M. Image-based food portion size estimation using a smartphone without a fiducial marker. Public Health Nutr., 1–13 (2018). doi:10.1017/S136898001800054X. https://www.cambridge.org/core/journals/public-health-nutrition/article/imagebased-food-portion-size-estimation-using-a-smartphone-without-a-fiducial-marker/47ED461DDE607FE0C7E6D70168E80BFA

10. Liao, H.-C., Lim, Z.-Y. & Lin, H.-W. Food intake estimation method using short-range depth camera. In Proc. 2016 IEEE Int. Conf. Signal and Image Process. (ICSIP), 198–204 (2016). doi:10.1109/SIPROCESS.2016.7888252. https://www.semanticscholar.org/paper/ed0cf9d78ffbc54be9c76e97a67d61a55b918a8c

11. Fang, S., Zhu, F., Jiang, C., Zhang, S., Boushey, C. J. & Delp, E. J. A comparison of food portion size estimation using geometric models and depth images. In Proc. 2016 IEEE Int. Conf. Image Process. (ICIP), 26–30 (2016). doi:10.1109/ICIP.2016.7532312. https://www.semanticscholar.org/paper/456c7833fcd432c4a0a959214e5c46535608a8f4

12. Moyen, A. et al. Relative Validation of an Artificial Intelligence-Enhanced, Image-Assisted Mobile App for Dietary Assessment in Adults: Randomized Crossover Study. J. Med. Internet Res. 24, e40449 (2022). doi:10.2196/40449. https://pubmed.ncbi.nlm.nih.gov/36409539/

13. Jefferson, K. et al. Rx Food App: A Proof-of-Concept Study of an Image-Based Dietary Assessment Mobile Application. Curr. Dev. Nutr. 5(Suppl 2), 1001 (2021). doi:10.1093/cdn/nzab052_004. https://pmc.ncbi.nlm.nih.gov/articles/PMC8180873/

14. Google. Gemini API pricing. (accessed 2026-01-29). https://ai.google.dev/gemini-api/docs/pricing

15. Fridolfsson, J., Sjoberg, E., Thiwang, M. & Pettersson, S. Performance Evaluation of 3 Large Language Models for Nutritional Content Estimation from Food Images. Curr. Dev. Nutr. 9, 107556 (2025). doi:10.1016/j.cdnut.2025.107556. https://doi.org/10.1016/j.cdnut.2025.107556

16. Tanabe, H. & Yanai, K. Reasoning-Driven Food Energy Estimation via Multimodal Large Language Models. Nutrients 17, 1128 (2025). doi:10.3390/nu17071128. https://pubmed.ncbi.nlm.nih.gov/40218886/

17. OpenAI. GPT-4 Technical Report. arXiv 2303.08774 (2023). https://arxiv.org/abs/2303.08774

18. Gemini Team. Gemini: A family of highly capable multimodal models. arXiv 2312.11805 (2023). https://arxiv.org/abs/2312.11805

19. Radford, A. et al. Learning Transferable Visual Models From Natural Language Supervision. arXiv 2103.00020 (2021). https://arxiv.org/abs/2103.00020

20. Alayrac, J.-B., et al. Flamingo: a Visual Language Model for Few-Shot Learning. arXiv 2204.14198 (2022). https://arxiv.org/abs/2204.14198

21. Chen, X., et al. PaLI: A Jointly-Scaled Multilingual Language-Image Model. arXiv 2209.06794 (2022). https://arxiv.org/abs/2209.06794

22. Li, J. et al. BLIP: Bootstrapping Language-Image Pre-training for Unified Vision-Language Understanding and Generation. arXiv 2201.12086 (2022). https://arxiv.org/abs/2201.12086

23. Li, J. et al. BLIP-2: Bootstrapping Language-Image Pre-training with Frozen Image Encoders and Large Language Models. arXiv 2301.12597 (2023). https://arxiv.org/abs/2301.12597

24. Thames, Q. et al. Nutrition5k: Towards automatic nutritional understanding of generic food. In Proc. IEEE/CVF Conf. Comput. Vis. Pattern Recognit. 8903–8911 (2021). https://openaccess.thecvf.com/content/CVPR2021/html/Thames_Nutrition5k_Towards_Automatic_Nutritional_Understanding_of_Generic_Food_CVPR_2021_paper.html

25. Qwen Team. Qwen2-VL: Enhancing vision-language model’s perception of the world at any resolution. arXiv 2409.12191 (2024). https://arxiv.org/abs/2409.12191

26. FatSecret. Platform API documentation. (accessed 2026-01-27). https://platform.fatsecret.com/api/

27. Wei, J. et al. Chain-of-Thought Prompting Elicits Reasoning in Large Language Models. arXiv 2201.11903 (2022). https://arxiv.org/abs/2201.11903

28. Yao, S. et al. ReAct: Synergizing Reasoning and Acting in Language Models. arXiv 2210.03629 (2022). https://arxiv.org/abs/2210.03629

29. Shinn, N. et al. Reflexion: Language Agents with Verbal Reinforcement Learning. arXiv 2303.11366 (2023). https://arxiv.org/abs/2303.11366

30. Lin, L. I. A concordance correlation coefficient to evaluate reproducibility. Biometrics 45, 255–268 (1989). https://pubmed.ncbi.nlm.nih.gov/2720055/

31. Lucassen, D. A., Willemsen, R. F., Geelen, A., Brouwer-Brolsma, E. M., & Feskens, E. J. M. The accuracy of portion size estimation using food images and textual descriptions of portion sizes: an evaluation study. Journal of Human Nutrition and Dietetics 34(6), 945–952 (2021). doi:10.1111/jhn.12878.

32. Madaan, A. et al. Self-Refine: Iterative Refinement with Self-Feedback. Advances in Neural Information Processing Systems 36 (NeurIPS 2023) (2023). doi:10.48550/arXiv.2303.17651.

33. Dhuliawala, S., Komeili, M., Xu, J., Raileanu, R., Li, X., Celikyilmaz, A. & Weston, J. Chain-of- Verification Reduces Hallucination in Large Language Models. In Findings of the Association for Computational Linguistics: ACL 2024, 3563–3578 (2024). doi:10.18653/v1/2024.findings-acl.212.

34. OpenAI. API pricing. (accessed 2026-01-29). https://openai.com/api/pricing

35. Khlaisamniang, N. et al. Decomposing Food Images for Better Nutrition Analysis: a Nutritionist-Inspired Two-Step Multimodal LLM Approach. In Proc. IEEE/CVF Conf. Comput. Vis. Pattern Recognit. Workshops (CVPRW) (2025). doi:10.1109/CVPRW67362.2025.00053. https://www.semanticscholar.org/paper/5bfdab159345a3ac5dd72681fdc3e9f8bc0662c7

36. Cinar, E. N., Ozler, E., Arslan, S. & Yilmaz, S. Image-based nutritional assessment: Evaluating the performance of ChatGPT-4o on simple and complex meals. J. Food Compos. Anal. (2026). doi:10.1016/j.jfca.2025.108843. https://doi.org/10.1016/j.jfca.2025.108843

37. Ma, Z. et al. Food-500Cap: A Fine-Grained Food Caption Benchmark for Evaluating Vision- Language Models. arXiv 2308.03151 (2023). https://arxiv.org/abs/2308.03151

